# SARS-CoV-2 Alpha, Beta and Delta variants display enhanced Spike-mediated Syncytia Formation

**DOI:** 10.1101/2021.06.11.448011

**Authors:** Maaran Michael Rajah, Mathieu Hubert, Elodie Bishop, Nell Saunders, Remy Robinot, Ludivine Grzelak, Delphine Planas, Jérémy Dufloo, Stacy Gellenoncourt, Alice Bongers, Marija Zivaljic, Cyril Planchais, Florence Guivel-Benhassine, Françoise Porrot, Hugo Mouquet, Lisa Chakrabarti, Julian Buchrieser, Olivier Schwartz

**Author notes:** Second co-authors (Mathieu Hubert, Elodie Bishop, Nell Saunders). Last co-authors (Julian Buchrieser, Olivier Schwartz).

## Abstract

Severe COVID-19 is characterized by lung abnormalities, including the presence of syncytial pneumocytes. Syncytia form when SARS-CoV-2 spike protein expressed on the surface of infected cells interacts with the ACE2 receptor on neighbouring cells. The syncytia forming potential of spike variant proteins remain poorly characterized. Here, we first assessed Alpha and Beta spread and fusion in cell cultures. Alpha and Beta replicated similarly to D614G reference strain in Vero, Caco-2, Calu-3 and primary airway cells. However, Alpha and Beta formed larger and more numerous syncytia. Alpha, Beta and D614G fusion was similarly inhibited by interferon induced transmembrane proteins (IFITMs). Individual mutations present in Alpha and Beta spikes differentially modified fusogenicity, binding to ACE2 and recognition by monoclonal antibodies. We further show that Delta spike also triggers faster fusion relative to D614G. Thus, SARS-CoV-2 emerging variants display enhanced syncytia formation.

**Synopsis:** The Spike protein of the novel SARS-CoV-2 variants are comparative more fusogenic than the earlier strains. The mutations in the variant spike protein differential modulate syncytia formation, ACE2 binding, and antibody escape.

- The spike protein of Alpha, Beta and Delta, in the absence of other viral proteins, induce more syncytia than D614G
- The ACE2 affinity of the variant spike proteins correlates to their fusogenicity
- Variant associated mutations P681H, D1118H, and D215G augment cell-cell fusion, while antibody escape mutation E484K, K417N and Δ242-244 hamper it.
- Variant spike-mediated syncytia formation is effectively restricted by IFITMs

## Introduction

SARS-CoV-2 was initially discovered during an outbreak in Wuhan, China, before it became pandemic (Huang *et al*, 2020a). Since its emergence, the ancestral Wuhan strain has been supplanted by variants harboring a variety of mutations. Several of these mutations occur in the highly antigenic Spike (S) protein which endowed many of the variants with the ability to evade part of the neutralizing antibody response (Liu *et al*, 2021c; Planas *et al*, 2021a; Rees-Spear *et al*, 2021; Starr *et al*, 2021; Weisblum *et al*, 2020). Individual amino-acid changes in the S protein also affect viral fitness. One of the earliest identified variants contained the D614G mutation in S protein, which increased infectivity without significantly altering antibody neutralization (Yurkovetskiy *et al*, 2020). Several other variants have since emerged and have become globally dominant, including Alpha (B.1.1.7) first identified in the United Kingdom, Beta (B.1.351) identified in South Africa, Gamma (P.1 & P.2) identified in Brazil, and Delta (B.1.617.2) identified in India (Buss *et al*, 2021; Frampton *et al*, 2021; Planas *et al*, 2021b; Sabino *et al*, 2021; Tegally *et al*, 2020; Yadav *et al*, 2021). Some variants are more transmissible but their impact on disease severity is debated (Davies *et al*, 2021; Kemp *et al*, 2021; Korber *et al*, 2020).

Clinically, SARS-CoV-2 infections range from asymptomatic or febrile respiratory disorders to severe lung injury characterized by vascular thrombosis and alveolar damage (Bussani *et al*, 2020). The deterioration of respiratory tissue is likely a result of both virus-induced cytopathicity and indirect immune-mediated damage (Buchrieser *et al*, 2020; Zhang *et al*, 2020; Zhou *et al*, 2020; Zhu *et al*, 2020). A peculiar dysmorphic cellular feature is the presence of large infected multinucleated syncytia; predominately comprised of pneumocytes (Braga *et al*, 2021; Bussani *et al*., 2020; Sanders *et al*, 2021). Other coronaviruses including SARS-CoV-1, MERS-CoV, and HKU1 also induce syncytia formation in patient tissues and cell culture systems (Chan *et al*, 2013; Dominguez *et al*, 2013; Franks *et al*, 2003; Qian *et al*, 2013). Syncytial cells may compound SARS-CoV-2 induced cytopathicity, play a role in viral persistence and dissemination and could be a pathological substrate for respiratory tissue damage (Braga *et al*., 2021; Buchrieser *et al*., 2020; Sanders *et al*., 2021). Release of syncytial cells may contribute to the overall infectious dose (Beucher *et al*, 2021). Heterocellular syncytia containing lymphocytes have also been documented in the lungs of infected patients (Zhang *et al*, 2021).

The SARS-CoV-2 S protein is a viral fusogen. The interaction of trimeric S with the ACE2 receptor and its subsequent cleavage and priming by surface and endosomal proteases results in virus-cell fusion (Hoffmann *et al*, 2020). Merging of viral and cellular membranes allows for viral contents to be deposited into the cell to begin the viral life cycle. Within the cell, newly synthesized spike, envelope and membrane proteins are inserted into the endoplasmic reticulum (ER), and trafficked and processed through the ER-Golgi network (Cattin-Ortolá *et al*, 2021; Duan *et al*, 2020; Nal *et al*, 2005). Virion are formed by budding into ER-Golgi membranes and are then transported to the surface in order to be released from the cell (Klein *et al*, 2020). While the majority of the S protein is sequestered within the ER, motifs within its cytoplasmic tail allow for leakage from the Golgi apparatus and localization at the plasma membrane (Cattin-Ortolá *et al*., 2021). The S protein at the surface of an infected cell interacts with receptors on adjacent cells, fusing the plasma membranes together and merging the cytoplasmic contents. We and others had previously shown that the S protein interacting with the ACE2 receptor induces cell-cell fusion (Braga *et al*., 2021; Buchrieser *et al*., 2020; Lin *et al*, 2021; Sanders *et al*., 2021; Zhang *et al*., 2021).The TMPRSS2 protease further augments cell-cell fusion (Barrett *et al*, 2021; Buchrieser *et al*., 2020; Hornich *et al*, 2021).

The S protein is comprised of S1 and S2 subunits. The S1 subunit includes the N-terminal domain (NTD) and the receptor binding domain (RBD). The function of the NTD has yet to be fully elucidated but it may be associated with glycan binding, receptor recognition and pre-fusion-to-post fusion conformational changes. The NTD is also targeted by neutralizing antibodies (Chi *et al*, 2020; Krempl *et al*, 1997; Zhou *et al*, 2019). The RBD interacts with the ACE2 receptor and is the main target for neutralizing antibodies (Huang *et al*, 2020b). The S2 domain consists of the fusion peptide (FP), heptapeptide repeat sequences 1 and 2, (HR1 and HR2), the transmembrane anchor, and the C-terminal domain. The FP inserts into the target membrane by disrupting the lipid bilayer and anchors the target membrane to the fusion machinery (Huang *et al*., 2020b). This exposes regions of HR1 that interact with HR2, forming a flexible loop that brings the membranes together to facilitate fusion (Huang *et al*., 2020b). The versatility of the S protein suggests that any mutations that have arisen are of particular concern as they can affect fusogenicity, antibody recognition, affinity to ACE2, proteolytic processing and incorporation into virions. There is a general paucity of information regarding how the mutations associated with variant S proteins contribute to cell-cell fusion.

S-mediated cell-cell fusion is sensitive to innate immunity components. The interferon response to SARS-CoV-2 is one of the key factors down-modulating viral entry and replication, and deficiencies in the interferon response are associated with severe or critical COVID-19 (Arunachalam *et al*, 2020; Bastard *et al*, 2021; Bastard *et al*, 2020; Hadjadj *et al*, 2020; van der Made *et al*, 2020). SARS-CoV-2 induced syncytia formation by the Wuhan strain is restricted by innate immunity, in part through the action of interferon induced transmembrane proteins (IFITMs) (Buchrieser *et al*., 2020). IFITM1, 2 and 3 are restriction factors which display antiviral activity against a variety of enveloped viruses including SARS-CoV-2; likely by increasing membrane rigidity and hindering virus-cell fusion (Shi *et al*, 2021). Their effectiveness at restricting cell-cell fusion of novel variants has yet to be assessed.

Here, we compared the replication and syncytia forming potential of D614G, Alpha and Beta viruses in human cell lines and primary airway cells. We further characterized the fusogenicity of the Alpha and Beta variant S proteins and the individual contribution of each of the component mutations in syncytia formation, ACE2 binding and evasion from a panel of antibodies. Finally, we examined the syncytia forming potential and ACE2 binding capacity of the novel Delta variant spike.

## Results

### Comparative Replication Kinetics of SARS-CoV-2 Variants

We compared the replication kinetics of SARS-CoV-2 variants in relevant cell cultures. We first infected Caco-2, Calu-3 and Vero cells with Alpha, Beta, and D614G variants and generated multistep growth curves (Fig. 1). Cell were infected at an equivalent, non-saturating MOI, initially titrated in Vero cells (Fig. EV1A). Viral replication was assessed at 24, 48, and 72 hours by flow cytometry upon staining with the pan-SARS-CoV-2 anti-S mAb102 human monoclonal antibody (Planas *et al*., 2021a) and then gating for S+ cells (Fig. EV1B). Globally, the variants replicated similarly (Fig. 1). This similar replication was observed at different MOIs (Fig. EV1A). There were subtle differences at 24h post-infection, depending on the cell line and the variant. For instance, Beta replicated slightly more than D614G in Caco-2 cells whereas Alpha replicated slight less than D614G in Vero cells (Fig. 1A, C). Viral release at each time point was also assessed by extracting RNA from the supernatant and performing RT-qPCR for the gene encoding the N protein. Viral release was again roughly similar with the different variants, especially at early time points. Alpha produced moderately more virus than D614G in all cell lines at later time points (Fig. 1A-C). Beta produced more virus than D614G in Caco-2 but less in Calu-3 cells at later time points (Fig. 1A, B).

**Figure 1.**
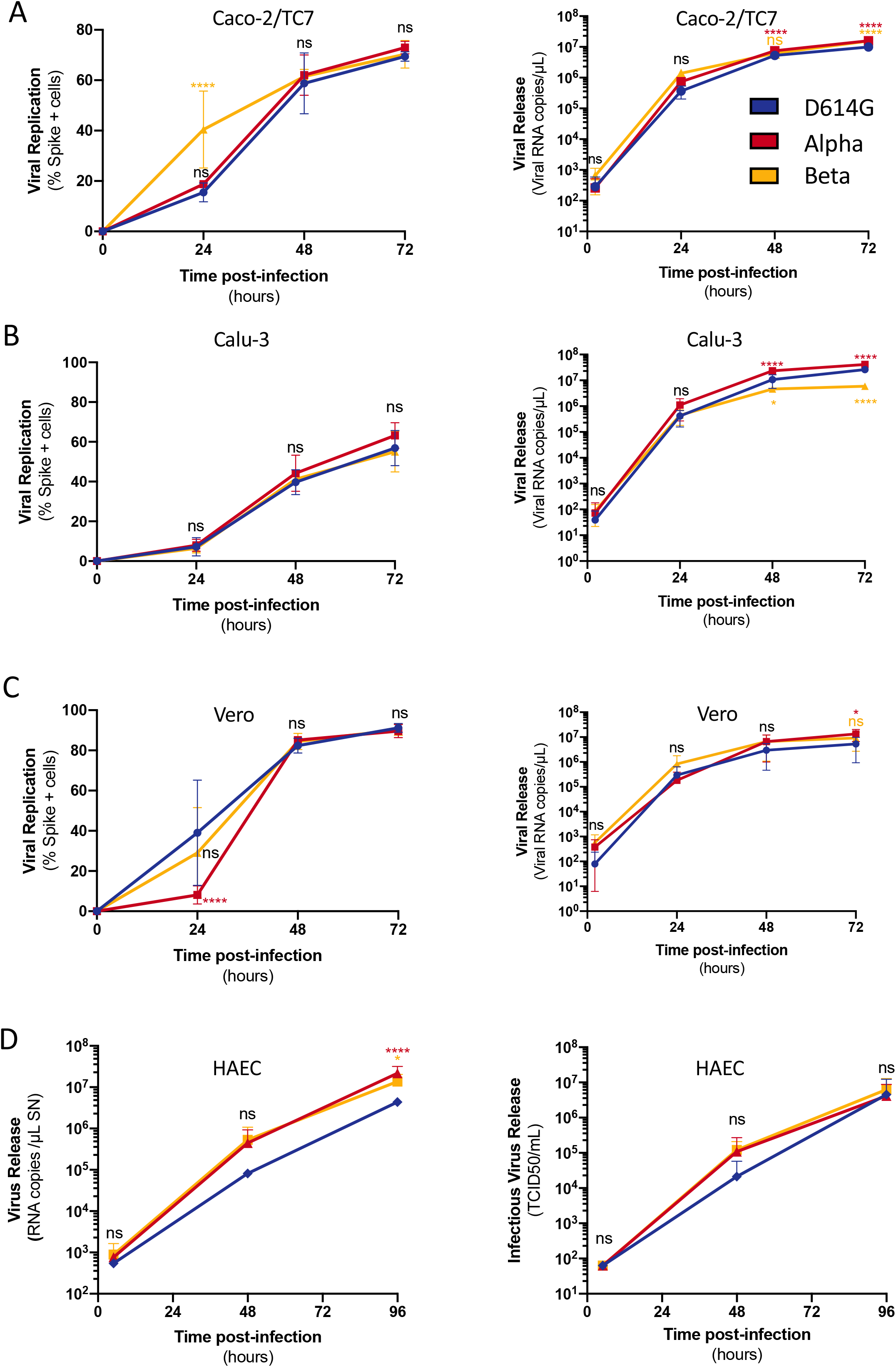
Replication kinetics of D614G, Alpha and Beta variants in cell culture. Cells were infected at the indicated MOI. Viral replication (Left) and release (Right) were assessed by flow cytometry and RT-qPCR. **A)** Caco2/TC7 cells (MOI 0.01) B) Calu-3 cells (MOI 0.001) **C)** Vero cells (MOI 0,01) **D)** primary human airway epithelial cells (HAEC) virus release (Right) and infectious virus release (Left) (MOI 0.01). Data are mean ± SD of at least 3 independent experiments. Statistical analysis: Mixed-effect analysis or Two-way ANOVA compared to D614G reference, ns: non-significant, *P < 0. 05, **P < 0.01, ***P < 0.001, ****P < 0.0001.

We then used the MucilAirB^TM^ model, which consists of primary human airway epithelial cells (HAEC) grown over a porous membrane and differentiated at the air-liquid interface for over 4 weeks. This relevant model is susceptible to SARS-CoV-2 infection (Pizzorno *et al*, 2020; Robinot *et al*, 2020; Robinot *et al*, 2021; Touret *et al*, 2021). The cells were infected with each variant at a similar low viral inoculum (10^4^ TCID50). Viral RNA and infectious virus release were monitored over 96h by RT-qPCR and TCID50. Alpha and Beta variants produced slightly more extracellular viral RNA than D614G at later time points but not significantly higher levels of infectious particles (Fig. 1D).

Taken together our data show that Alpha and Beta variants replicate similarly to the ancestral D614G strain in a panel of human cell lines and in primary cells, with some slight differences.

### Syncytia formation in cells infected with SARS-CoV-2 variants

We next assessed the potential of SARS-CoV-2 variants to induce syncytia. In order to visualize cell-cell fusion, we employed our previously described S-Fuse assay, using U2OS-ACE2 GFP-split cells (Buchrieser *et al*., 2020). In the GFP-split complementation system, two cell lines containing half of the reporter protein are co-cultured, producing a GFP signal only upon fusion (Fig. 2A). Upon infection of S-Fuse cells, we noticed that the Alpha and Beta variants formed larger and more numerous infected syncytia than either D614G or the ancestral Wuhan strain (Fig. EV2A). We then characterized quantitatively the differences in fusogenicity by calculating the total syncytia (GFP) area and then normalizing it to nuclei number (Hoechst) (Fig. EV2B). Relative to D614G, Alpha and Beta variants produced significantly more syncytia, approximately 4.5 and 3-fold respectively, after 20h of infection with the same MOI (Fig. 2B **and** EV3A). In order to characterize syncytia formation in a cell line expressing endogenous ACE2, we generated Vero cells carrying the GFP-split system. After 48h of infection with the same MOI, we again found that Alpha and Beta variants produced significantly more syncytia than D614G (Fig. 2C **and** EV3B) despite similar infection levels (Fig. 1C). Of note, D614G produced similar levels of syncytia as the Wuhan strain in both Vero and S-Fuse cells (Fig. 2 **and** EV3A-B).

**Figure 2.**
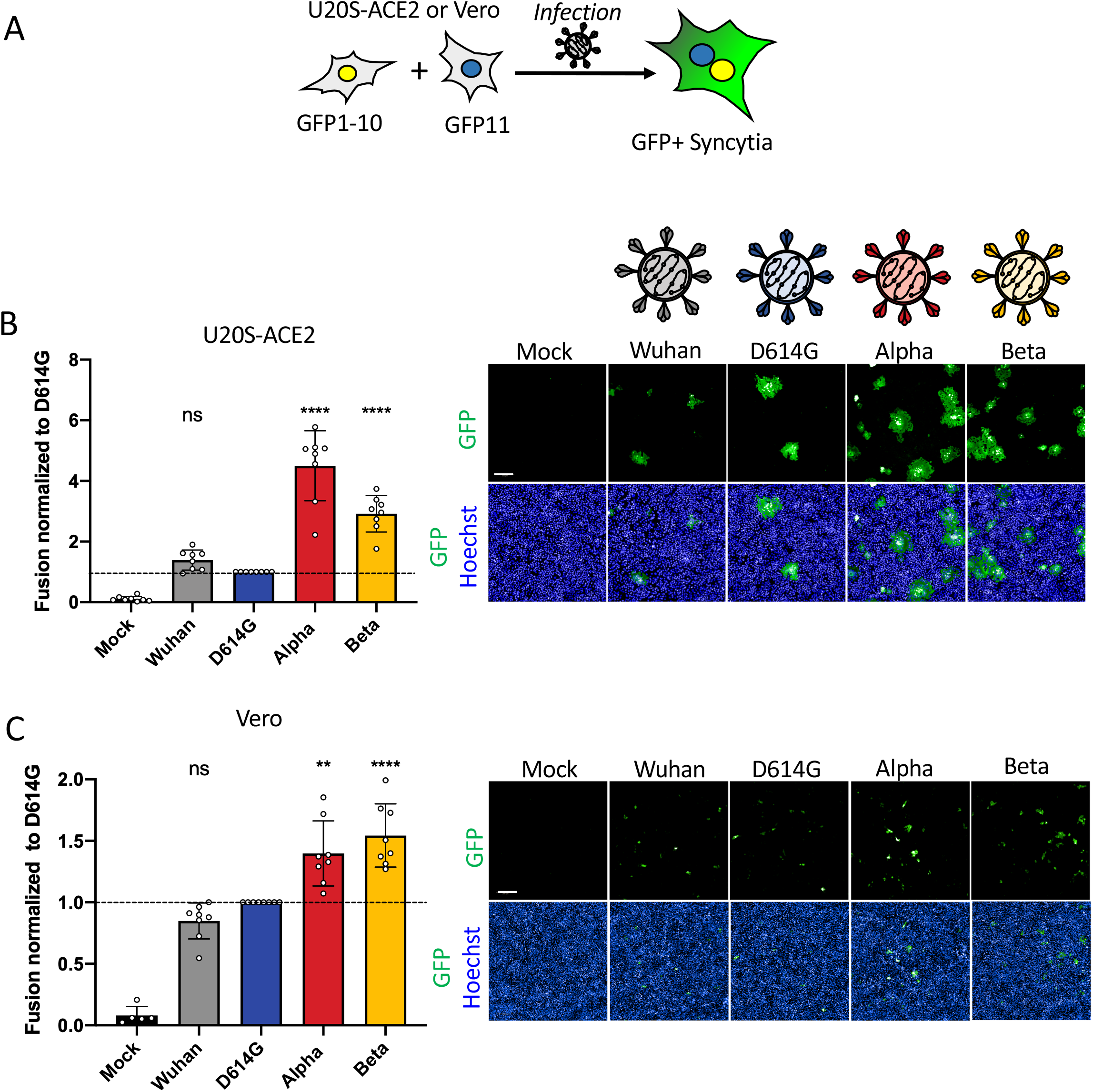
SARS-CoV-2 variant infection increases formation of syncytia in U2OS-ACE2 and Vero GFP-split cells. **(A)** U2OS-ACE2 or Vero cells expressing either GFP 1-10 or GFP 11 (1:1 ratio) were infected 24h after plating and imaged 20h (U2OS-ACE2) or 48h (Vero) post-infection. **(B) Left Panel:** Fusion was quantified by GFP area/ number of nuclei and normalized to D614G for U2OS-ACE2 20h post infection at MOI 0.001. **Right Panel:** Representative images of U2OS-ACE2 20h post infection, GFP-Split (Green) and Hoechst (Blue). Top and bottom are the same images with and without Hoechst channel. **(C) Left Panel:** Quantified fusion of Vero cells infected at MOI 0.01. **Right Panel:** Representative images of Vero cells 48h post infection, GFP-Split (Green) and Hoechst (Blue). Scale bars: 200 µm. Data are mean ± SD of 8 independent experiments. Statistical analysis: One-way ANOVA compared to D614G reference, ns: non-significant, *P < 0. 05, **P < 0.01, ***P < 0.001, ****P < 0.0001.

Therefore, Alpha and Beta variants appear more fusogenic than D614G in S-Fuse and Vero cells.

### Syncytia formation in cells expressing variant Spikes

Since syncytia formation is a consequence of the S protein expressed on the surface of an infected cell interacting with ACE2 receptors on neighboring cell, we sought to compare the fusogenic potential of the individual variant S proteins. We introduced the D614G mutation into the Wuhan protein and designed plasmids to express Alpha and Beta S proteins. We transfected the respective plasmids into Vero GFP-split cells and quantified syncytia formation 18h later (Fig. 3A). Alpha and Beta S proteins were 2 and 1.7-fold more fusogenic than D614G S, respectively (Fig. 3B). The Wuhan S was slightly less fusogenic that the D614G S (Fig. 3B). We then verified that the variation in S mediated fusion was not due to differential cell surface levels. We transfected 293T cells, which lack ACE2 and thus do not fuse upon S expression, with the different variant plasmids in order to assess S protein surface levels by flow cytometry. The variants S proteins were equally expressed after transfection (Fig. EV4A-C).

**Figure 3.**
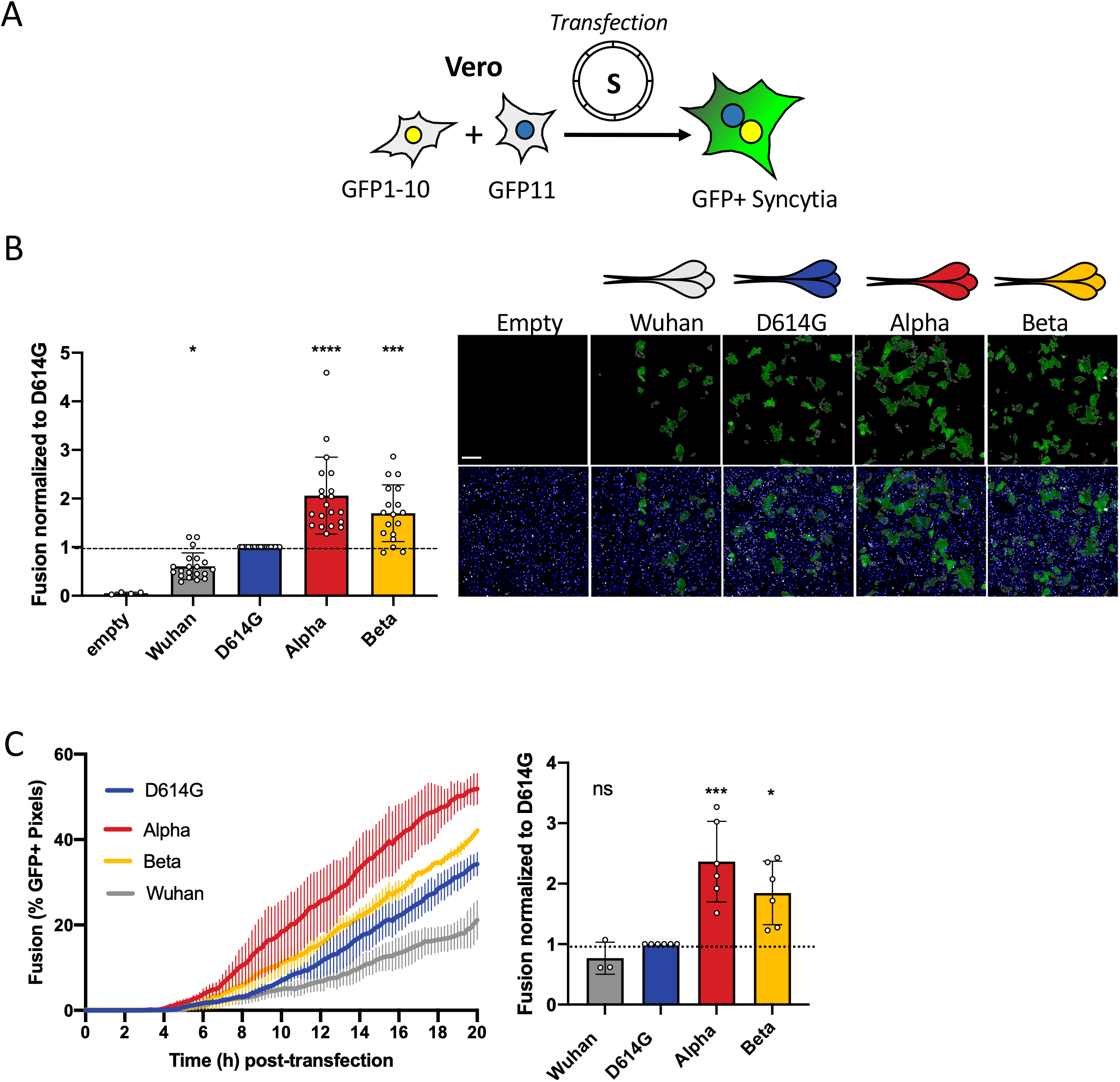
Alpha and Beta SARS-CoV-2 S proteins induce more robust syncytia formation than D614G. **(A)** Vero GFP-split cells were transfected with variant S proteins and imaged 18h post-transfection. **(B) Left Panel**: Fusion was quantified by GFP area/ number of nuclei and normalized to D614G for each of the transfected variant spike proteins. **Right Panel**: Representative images of Vero GFP-split cells 18h post-transfection, GFP (Green) and Hoechst (Blue). Top and bottom are the same images with and without Hoechst channel. **(C) Left Panel**: Quantification of variant S protein mediated fusion in Vero-GFP split cells by video microscopy. Results are mean ± SD from three fields per condition from one representative experiment. **Right Panel**: Fusion quantification of at least 3 independent video microscopy experiments, 20h post transfection, normalized to D614G. Scale bars: 200 µm. Data are mean ± SD of at least 3 independent experiments. Statistical analysis: One-way ANOVA compared to D614G reference, ns: non-significant, *P < 0. 05, **P < 0.01, ***P < 0.001, ****P < 0.0001.

We then measured the kinetics of syncytia formation induced by the different S proteins in Vero GFP-Split cells. We conducted a comparative video-microscopy analysis where cell-cell fusion could be visualized as soon as 6h post-transfection. The fusion kinetics of Alpha S protein was more rapid than any of the other variants (Fig. 3C **and Movie E1)**. Beta also induced significantly faster fusion than D614G, whereas the Wuhan S was the slowest of all the compared proteins (Fig. 3C **and Movie E1)**.

We then asked whether the TMPRSS2 protease, that cleaves S and facilitate viral fusion, may act differently on the variant S proteins. To this aim, we expressed the different variant S proteins without or with TMPRSS2 in 293T cells. We examined the processing of the different S by western blot and the surface levels by flow cytometry. The cleavage profile induced by TMPRSS2 and the surface levels of the different variant S proteins were similar (Fig EV4E).

Altogether, our data indicate that the S proteins of Alpha and Beta variants form more syncytia than D614G or Wuhan strains.

### Restriction of S mediated syncytia formation by IFN-β1 and IFITMs

As the variants did not show any major difference in replication under basal conditions, we next investigated if they were differently sensitive to the interferon response. To this aim, we pre-treated Vero cells or U2OS-ACE2 (S-Fuse) cells with increasing doses of IFN-β1 and infected them with the different variants. IFN-β1 was equally effective at reducing viral replication of D614G, Alpha, and Beta variants in Vero cells (Fig. EV5A). Preincubation of S-Fuse cell with IFN-β1 also abrogated infection and syncytia formation to the same extent with the different variants (Fig. EV5B). Therefore, IFN-β1 similarly inhibited viral replication and reduced syncytia formation by D614G, Alpha and Beta variants.

IFITMs are interferon stimulated transmembrane proteins that restrict early stages of the viral life cycle by inhibiting virus-cell fusion; likely by modifying the rigidity or curvature of membranes (Compton *et al*, 2014; Shi *et al*, 2017; Zani & Yount, 2018). IFITM1 localizes at the plasma membrane while IFITM2 and 3 transit through surface and localize in endo-lysosomal compartments (Buchrieser *et al*., 2020). We previously reported that IFITMs restrict Wuhan S mediated cell-cell fusion and that their activity was counteracted by the TMPRSS2 protease (Buchrieser *et al*., 2020). As infection with Alpha and Beta induce more syncytia, we further investigated if this resulted in an increased resistance to IFITM restriction. We thus characterized the impact of IFITMs on syncytia formed upon expression of D614G, Alpha and Beta S proteins in 293T cells. The variants were effectively restricted by IFITMs (Fig. EV5C-G). Of note, the three IFITMs were expressed at similar levels (not shown). The presence of TMPRSS2 increased fusion of all S proteins and reverted the restriction by IFITMs (Fig. EV5C-G). Taken together, our data show that Alpha and Beta variants induce more syncytia, but their S proteins remain similarly sensitive to IFITMs.

### Contribution of individual variant-associated mutations on Spike fusogenicity

We next sought to determine the contribution of each variant-associated mutation to cell-cell fusion. Both Alpha and Beta S proteins contain the N501Y mutation in the RBD and the D614G mutation in the S1/S2 cleavage site (Fig. 4A). Alpha S contains the Δ69/70 and ΔY144 deletions in the N-terminal domain (NTD), P681H and T716I mutations in the S1/S2 cleavage site, the S982A mutation in the heptad repeat 1 (HR1) site and the D1118H mutation in between HR1 and HR2. The Beta S is comprised of the L18F, D80A, D215G and Δ242-244 mutations in the NTD, K417N and E484K mutations in the receptor binding domain (RBD), and A701V in the S1/S2 cleavage site. We introduced individual mutations into the D614G background. Following reports of the emergence of the E484K mutation within the Alpha variant (Collier *et al*, 2021), we also generated a mutant Alpha S protein with the E484K mutation. We observed by flow cytometry that the mutant S proteins were similarly expressed at the cell surface (Fig. EV4A-C **and** EV6 A-D). We expressed each mutant into Vero GFP split and measured their potential to induce cell-cell fusion in comparison to D614G S protein.

**Figure 4.**
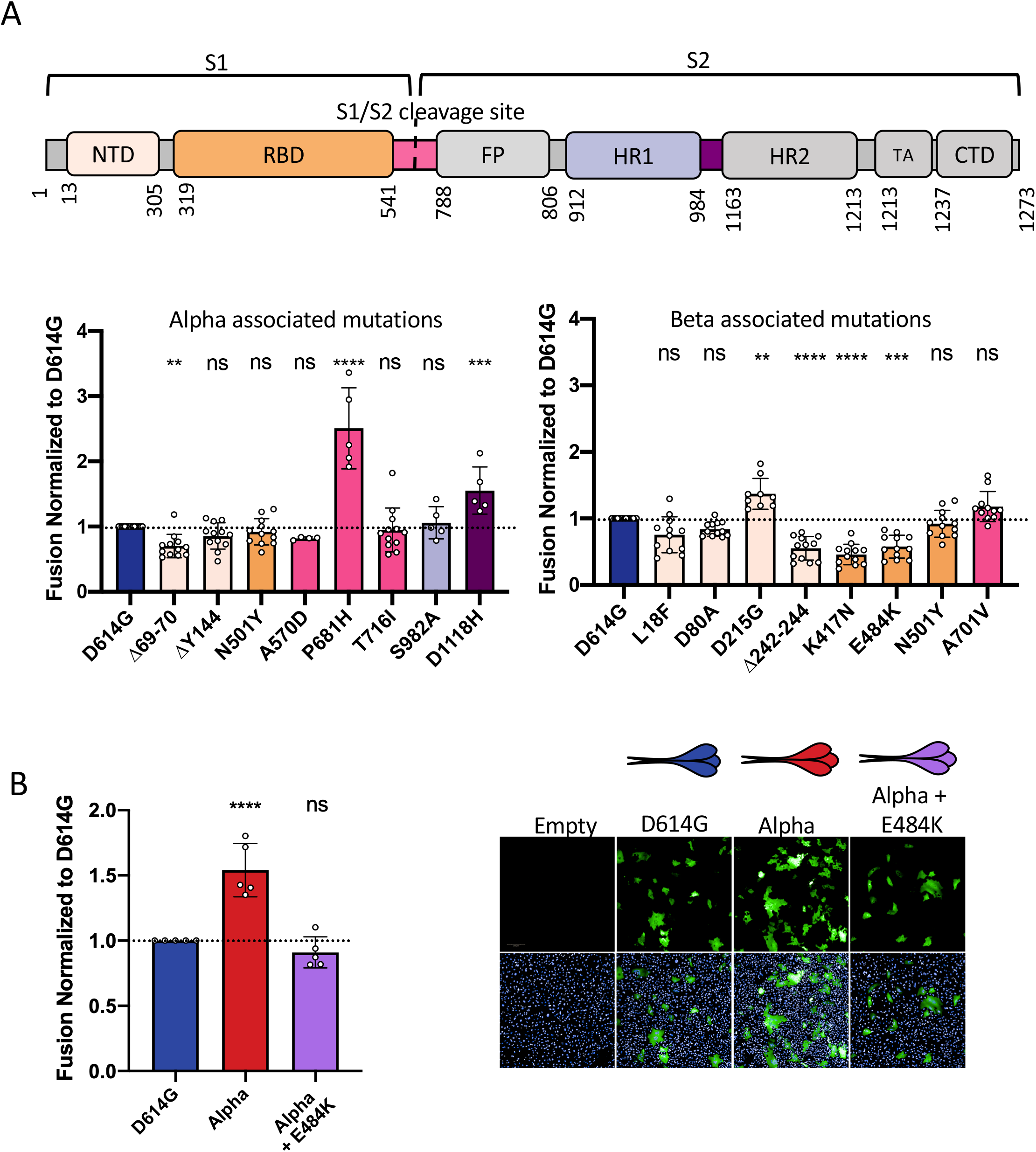
Mutations associated with Alpha and Beta S proteins differentially affect cell-cell fusion. **(A)** Schematic representation of the S protein colour coded for the functional regions: N-terminal domain (NTD), receptor binding domain (RBD), fusion peptide (FP), heptad repeat 1,2 (HR1, HR2), transmembrane anchor (TA), C-terminal domain (CTD). **(B) Left Panel:** Vero GFP-split cells were transfected with spike plasmids containing each of the individual mutations associated with Alpha variant in the D614G background. The amount of fusion was quantified at 20h and normalized to D614G reference plasmid. Colour code of each mutation corresponds to spike functional regions represented in (A). **Right Panel:** Quantified fusion for each of the individual spike mutations associated with Beta. Data set for N501Y and D614G reference mutations are duplicated between left and right panels for presentation as mutations are common to both variants. **(C) Left Panel:** Quantified fusion of the Alpha + E484K variant S protein normalized to D614G S. **Right Panel:** Representative images of fusion at 20h. Scale bar: 200 µm. Data are mean ± SD of at least 4 independent experiments. Top and bottom are the same images with and without Hoechst channel. Statistical analysis: statistics for both left and right panels of A were conducted together. One-way ANOVA compared to D614G reference, ns: non-significant, *P < 0. 05, **P < 0.01, ***P < 0.001, ****P < 0.0001.

Of the mutations that are associated with Alpha, we found that the Δ69/70 deletion in the RBD decreased cell-cell fusion whereas P681H and D1118H substitutions both increase fusion (Fig. 4A **and** EV6G). P681H displayed the greatest fusogenicity of all investigated mutations, being almost 2.5-fold higher than D614G S (Fig. 4A **and** EV6G). As previously mentioned, the introduction of the D614G mutation in the S1/S2 border of the Wuhan S protein also relatively increased fusion, stressing the importance of this cleavage site in fusogenicity (Fig. 3B).

Among the mutations associated with Beta, the Δ242-244 deletion, as well as K417N and E484K mutations in the RBD significantly decreased syncytia formation (Fig. 4A **and** EV6H). Only the D251G mutation in the NTD modestly increased syncytia formation relative to D614G (Fig. 4A **and** EV6H). The introduction of the E484K RBD mutation into the Alpha S protein significantly decreased its potential to form syncytia, despite not changing cell surface expression, further supporting the mutation’s restrictive effect on cell-cell fusion (Fig. 4B **and** EV4B). Taken together, our data suggests that variant S proteins are comprised of mutations that play contrasting roles in cell-cell fusion. P681H, D1118H, and D215G substitutions facilitate fusion, whereas mutations Δ69/70, Δ242-244, K417N, and E484K antagonize cell-cell fusion.

### Binding of S proteins bearing individual variant-associated mutations to ACE2

We next explored the impact of variant-associated mutations on S binding to the ACE2 receptor. To this aim, we transiently expressed each mutant protein in 293T cells. Cells were then stained with a serial dilution of soluble biotinylated ACE2, revealed with fluorescent streptavidin and then analyzed by flow cytometry (Fig. 5A). Titration binding curves were generated and EC50 (the amount of ACE2 needed for 50% binding) was calculated. The S protein of Alpha had the highest affinity to ACE2, confirming previous results by us and others (Planas *et al*., 2021)(Ramanathan *et al*, 2021). Alpha was sequentially followed by Beta, D614G, and Wuhan S (Fig. 5B **and** EV7A). As expected, mutations within the RBD had the most significant impact on ACE2 binding. N501Y found in both Alpha and Beta drastically increased ACE2 binding, in line with previous reports indicating that this mutation enhances affinity of the viral protein to its receptor (Ali *et al*, 2021; Luan *et al*, 2021; Tian *et al*, 2021). The K417N substitution present in the Beta S decreased ACE2 binding (Fig. 5B **and** EV7C). The E484K mutant had a slightly, but not significantly, higher binding to ACE2 (Fig. EV7C). This was corroborated by the observation that addition of the E484K mutation to Alpha S protein also slightly increased ACE2 binding (Fig. 5B **and** EV7A). Mutation in the S1/S2 cleavage site, HR1/HR2 sites or NTD did not have any significant impact on ACE2 binding (Fig. 5B **and** EV7B-E). It is worth noting that the NTD Δ242-244 mutant displayed a marginally lower binding to ACE2 (Fig. 5B **and** EV7B). Therefore, the N501Y mutation is the most significant contributor to increased ACE2 binding of the variants, though it does not affect cell-cell fusion on its own. The K417N, Δ242-244, and E484K mutations restrict fusogenicity but differently effect ACE2 binding; with the former two decreasing affinity and the latter slightly increasing.

**Figure 5.**
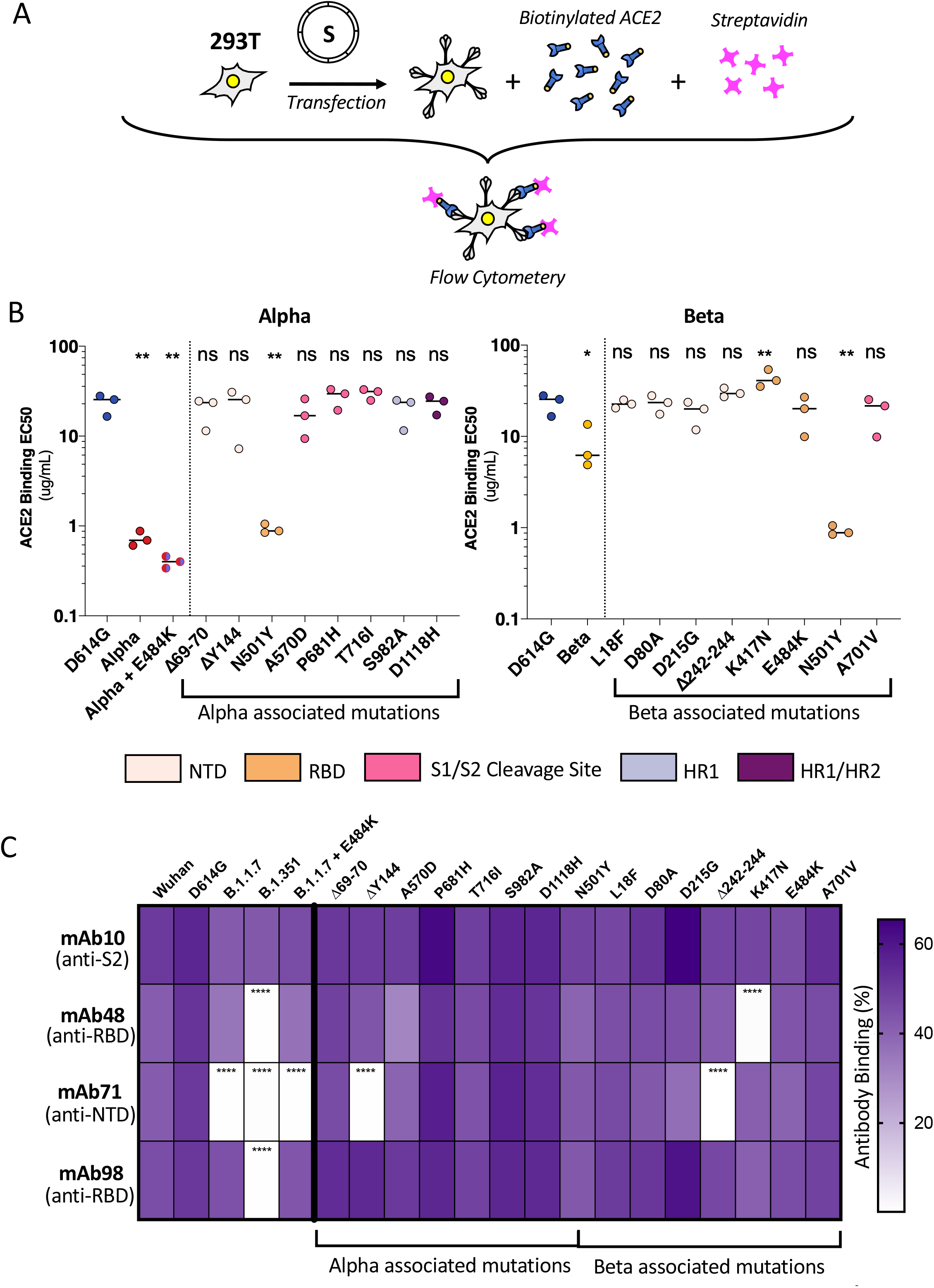
ACE2 and monoclonal antibody binding to S proteins with Alpha and Beta associated mutations. **(A)** 293T cells were transfected S proteins with each variant-associated mutation for 24h and stained with biotinylated ACE2 and fluorescent streptavidin before analysis by flow cytometry. **(B) Left Panel:** EC50 values (concentration of ACE2 needed for 50% binding) for Alpha and associated mutations. Colour code corresponds to location on spike functional domains and lower EC50 values signifies higher affinity to ACE2 binding. **Right Panel:** EC50 values for Beta and associated mutations. Data set for N501Y and D614G reference mutations are duplicated between left and right panels as mutations are common to both variants. **(C)** Spike transfected 293T cells were stained with human monoclonal antibodies targeting the S2 (mAb10), RBD (mAb48 and mAb98), and the NTD (mAb71). Cells were analyzed by flow cytometry. The percentage of positive cells is indicated. Data are mean of at least 3 independent experiments. Statistical analysis: One-way ANOVA compared to D614G reference, ns: non-significant, *P < 0. 05, **P < 0.01, ***P < 0.001, ****P < 0.0001.

Therefore, ACE2 binding and fusogenicity are two functions of the S protein that can be partially deconvoluted through individual mutations.

### Antibody binding to S proteins bearing individual variant-associated mutations

We had previously found that certain neutralizing antibodies differentially affect SARS-CoV-2 D614G, Alpha and Beta variants (Planas *et al*., 2021a). For instance, neutralizing monoclonal antibody 48 (mAb48) restricts D614G virus but not Alpha or Beta variants (Planas *et al*., 2021a). We sought to determine which mutations in variant S proteins contributed to the lack of recognition by the neutralizing antibodies. To this aim, we assessed by flow cytometry the binding of a panel of four human monoclonal antibodies (mAbs) to the different S mutants. As a control we used mAb10, a pan-coronavirus antibody that targets an unknown but conserved epitope within the S2 region (Planchais, manuscript in preparation). mAb10 equally recognized all variants and associated individual mutations (Fig. 5C). mAb48 and mAb98 target the RBD and mAb71 the NTD (Planas *et al*., 2021b)(Planchais, manuscript in preparation). mAb48 did not recognize the Beta variant, and more specifically did not bind to the K417N mutant (Fig. 5C). The mAb71 recognized neither Alpha nor Beta variants and did not bind to their respective NTD ΔY144 and Δ242-244 mutations. The K417N and Δ242-244 mutations were also responsible for decreasing S-mediated fusion, suggesting a tradeoff between antibody escape and fusion (Fig. 5C). mAb98 did not recognize the Beta variant. However, none of the associated mutations were specifically responsible for the lack of binding (Fig. 5C), suggesting a combined effect on the structure of the S protein that may affect antibody escape.

Therefore, several of the mutations found in the variants spike proteins are advantageous in terms of antibody escape despite slightly reducing the ability the proteins to fuse.

### Spike mediated syncytia formation by the Delta variant

With the emergence and rapid spread of the Delta variant, we sought to characterize its potential to form syncytia. We recently showed that the Delta variant induce large syncytia in S-Fuse cells (Planas *et al*., 2021b). We thus compared the fusogenicity of the Delta S protein to that of D614G and Alpha. We transiently expressed the three S proteins in Vero-GFP split cells. The Delta S protein triggered more cell-cell fusion than the D614G variant but was similar to the Alpha S protein (Fig 6A). The fusion kinetic of the Delta S was also similar to Alpha but more rapid than D614G (Fig 6B). We confirmed that the variant S proteins were equally expressed on the surface by transfecting them into non-fusogenic 293T cells and performing flow cytometry upon staining with the pan-SARS-CoV-2 mAb129 (Fig EV4D). We next examined the ACE2 binding potential of Delta S protein using our aforementioned soluble biotinylated ACE2. The Delta S protein has a higher binding capacity to ACE2 than the D614G S protein, but the binding was lower than the Alpha S protein (Fig 6C).

**Figure 6.**
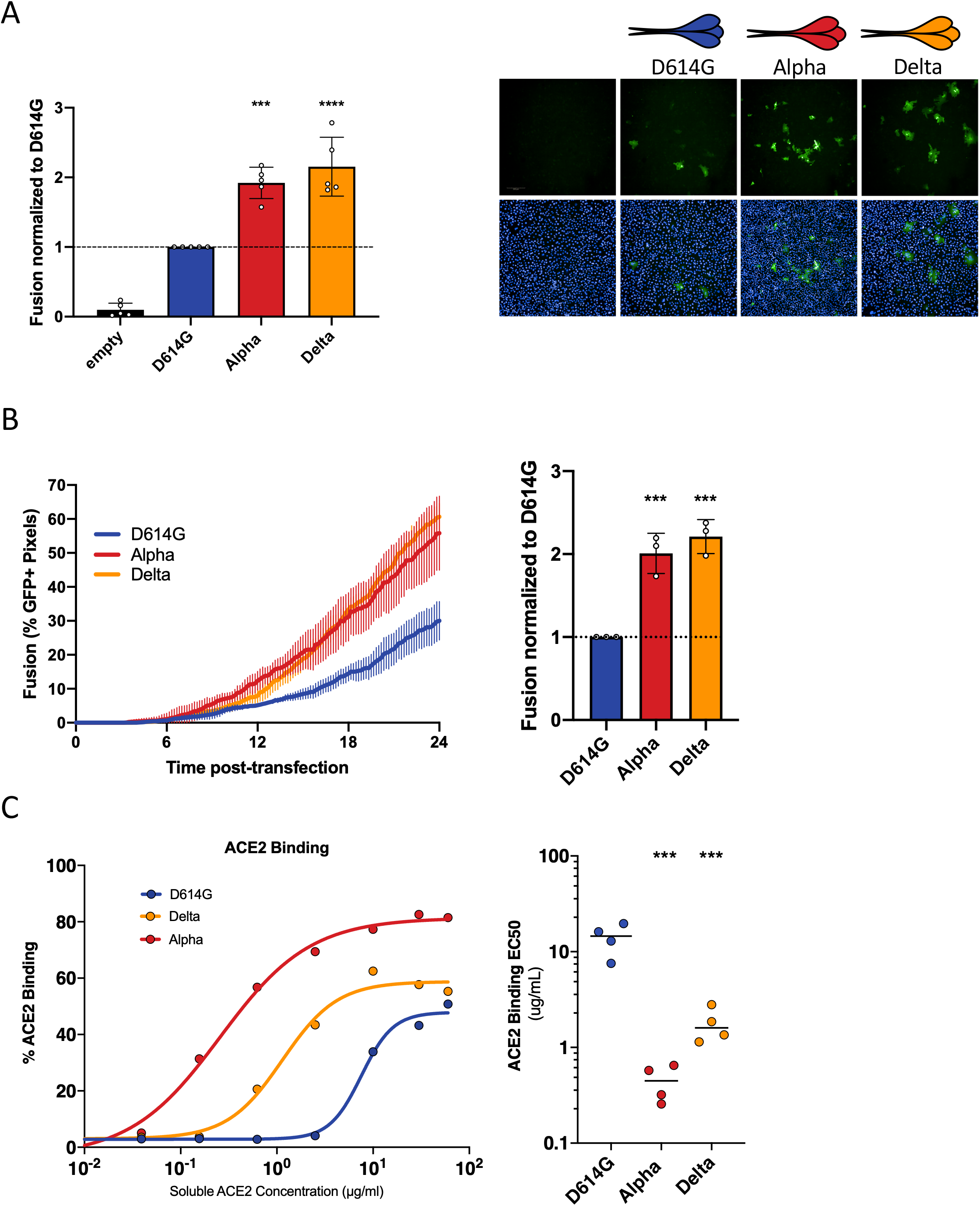
Delta SARS-CoV-2 S protein induces more syncytia formation and binds more to ACE2 than D614G. **(A)** Vero GFP-split cells were transfected with variant S proteins and imaged 18h post-transfection. **Left Panel**: Fusion was quantified by GFP area/ number of nuclei and normalized to D614G for each of the transfected variant spike proteins. **Right Panel**: Representative images of Vero GFP-split cells 18h post-transfection, GFP (Green) and Hoechst (Blue). Top and bottom are the same images with and without Hoechst channel. **(B) Left Panel**: Quantification of Delta S protein mediated fusion in Vero-GFP split cells by video microscopy. Results are mean ± SD from three fields per condition from one representative experiment. **Right Panel**: Fusion quantification of 3 independent video microscopy experiments, 20h post transfection, normalized to D614G. **(C) 293**T cells were transfected S proteins with each variant-associated mutation for 24h and stained with biotinylated ACE2 and fluorescent streptavidin before analysis by flow cytometry. **Left Panel:** Representative ACE2 binding dilution curves for the Delta variant in relation to Alpha and D614G. **Right Panel:** EC50 values (concentration of ACE2 needed for 50% binding) for Alpha for the Delta variant. Scale bars: 200 µm. Data are mean ± SD of at least 3 independent experiments. Statistical analysis: One-way ANOVA compared to D614G reference, ns: non-significant, *P < 0. 05, **P < 0.01, ***P < 0.001, ****P < 0.0001.

## Discussion

The replication and cytopathic effects of SARS-CoV-2 variants is under intense scrutiny, with contrasting results in the literature (Frampton *et al*., 2021; Hou *et al*, 2020; Leung *et al*, 2021; Liu *et al*, 2021b; Touret *et al*., 2021). For instance, there was no major difference in the replication kinetics of Alpha and D614G strains in some reports (Thorne *et al*, 2021; Touret *et al*., 2021), whereas others suggested that Alpha may outcompete D614G in a co-infection assay (Touret *et al*., 2021). Other studies proposed that the N501Y mutation may provide a replication advantage, whereas others suggested that N501Y is deleterious (Frampton *et al*., 2021; Hou *et al*., 2020; Leung *et al*., 2021; Liu *et al*., 2021b). These discrepant results may be due to the use of different experimental systems, viral strains, multiplicities of infection and cell types.

Here, we show that Alpha and Beta variants replicate to the same extent as the early D614G strain in different human cell lines and primary airway cells. Moreover, Alpha and Beta induced more cell-cell fusion than D614G. Increased fusion was observed in U2OS-ACE2 cells and in naturally permissive Vero cells. In agreement with infection data, transfection of Alpha and Beta S proteins, in the absence of any other viral factors, produced significantly more syncytia than D614G, which in turn, fused more than the Wuhan S. Comparative video microscopy analysis revealed that Alpha S fused the most rapidly, followed by Beta, D614G, and finally Wuhan. Thus, Alpha and Beta variants display enhanced S-mediated syncytia formation.

We further show that Alpha and Beta remain sensitive to restriction by IFN-β1. The fusion mediated by their respective S proteins is inhibited by IFITMs. This extends previous results by us and others demonstrating that ancestral Wuhan S is effectively inhibited by this family of restriction factors (Buchrieser *et al*., 2020; Shi *et al*., 2021). It has been recently reported in a pre-print that Alpha may lead to lower levels of IFN-β1 production by infected Calu-3 cells and may be less sensitive to IFN-β pre-treatment, when compared to first wave viral isolates (Thorne *et al*., 2021). We did not detect here differences of IFN-β1 sensitivity between the variants in Vero and U2OS-ACE2 cells. Again, these discrepant results may reflect inherent differences between Calu-3, Vero and U2OS-ACE2 cells, or the use of different viral isolates.

We then characterized the contribution of the individual mutations present in Alpha and Beta S proteins to their respective fusogenicity. The highly fusogenic Alpha S consists of more mutations that robustly increase fusion (P681H and D1118H) than mutations that decrease fusion (Δ69/70). In contrast, the Beta variant is comprised of several restrictive mutations (Δ242-244, K417N, and E484K) and only one mutation that modestly increased fusion (D215G). The strongest increase in fusion was elicited by the P681H mutation at the S1/S2 border. This mutation likely facilitates proteolytic cleavage of S and thus promotes S mediate cell-cell fusion. Indeed, the analogous P681R mutation present in B.1.617.2 and B.1.617.3 variants increases S1/S2 cleavage and facilitates syncytia formation (Ferreira *et al*, 2021; Jiang *et al*, 2020). Of note, another report with indirect assessment of variant S fusogenicity suggested a mild decrease or no difference in cell-cell fusion of Alpha and Beta relative to Wuhan S (Hoffmann *et al*, 2021). These previous experiments were performed in 293T cells at a late time-points (24 hours post-transfection), which may preclude detection of an accelerated fusion triggered by the variants.

We show that the binding of variant S to soluble ACE2 paralleled their fusogenicity. Alpha bound the most efficiently to ACE2, followed by Beta, D614G and finally Wuhan. However, the ACE2 affinity of S proteins carrying individual mutations did not exactly correlate to fusogenicity. For instance, the N501Y and D614G mutations drastically increased ACE2 affinity, but only D614G enhanced fusogenicity. The K417N substitution, and to a lesser degree Δ242-244, had a lower affinity to ACE2 and also restricted cell-cell fusion. The E484K mutation significantly restricts fusion, but mildly increases ACE2 affinity. This suggests that on the level of individual S mutations, the relationship between ACE2 affinity and increased fusogenicity is not always linear. Variant mutations may also confer advantages in an ACE2 independent manner. Indeed, recent work has suggested that the E484 mutation may facilitate viral entry into H522 lung cells, requiring surface heparan sulfates rather than ACE2 (Puray-Chavez *et al*, 2021). It would be of future interest to examine the syncytia formation potential of the variant mutations in other cell types.

We selected a panel of 4 mAbs that displayed different profiles of binding to Alpha, Beta, D614G and Wuhan S proteins. The mAb10 targeting the S2 domain recognized all variants and was used as a positive control. Wuhan and D614G were recognized by the three other antibodies, targeting either the NTD or RBD. Alpha lost recognition by the anti-NTD mAb71, whereas Beta was neither recognized by mab71 nor by the two anti-RBD antibodies mAb48 and mAb 98. Upon examining the potential of S proteins carrying individual mutations to bind to human monoclonal antibodies, we found that the ones that restrict (Δ242-244, K417N) or have no effect on fusogenicity (ΔY144) are also not recognized by some mAbs. This suggests that variant S proteins have undergone evolutionary trade off in some circumstances; selecting for mutations that provide antibody escape at the detriment of fusogenicity. In accordance with our findings, deep sequence binding analysis and in vitro evolution studies suggest the N501Y mutation increases affinity to ACE2 without disturbing antibody neutralization (Liu *et al*, 2021a; Starr *et al*., 2021; Zahradník *et al*, 2021). The E484K and K417N RBD mutations in the Beta variant may also increase ACE2 affinity, particularly when in conjunction with N501Y (Zahradník *et al*., 2021) (Nelson *et al*, 2021). However, the resulting conformational change of the S protein RBD may also decrease sensitivity to neutralizing antibodies (Nelson *et al*., 2021). Future work assessing the structural and conformational changes in the S protein elicited by a combination of individual mutations or deletions may further help elucidate the increased fusogenicity and antibody escape potential of the variants.

While we had previously shown that the interaction between the S protein on the plasma membrane with the ACE2 receptor on neighboring cells is sufficient to induce syncytia formation, there is compelling evidence of the importance of the TMPRSS2 protease in S activation (Buchrieser *et al*., 2020; Dittmar *et al*, 2021; Koch *et al*, 2021; Ou *et al*, 2021). We did not detect any major differences in the processing of the variant spike proteins by TMPRSS2. It will be worth further characterizing how the fusogenicity of variant associated mutations are influenced by other cellular proteases.

The presence of infected syncytial pneumocytes was documented in the lungs of patients with severe COVID-19 (Bussani *et al*., 2020; Tian *et al*, 2020; Xu *et al*, 2020). Syncytia formation may contribute to SARS-CoV-2 replication and spread, immune evasion and tissue damage. A report using reconstituted bronchial epithelia found that viral infection results in the formation and release of infected syncytia that contribute to the infectious dose (Beucher *et al*., 2021). The neutralizing antibody response to SARS-CoV-2 infection has divergent effect on cell-cell fusion, with some antibodies restricting S mediated fusion, while other increase syncytia formation (Asarnow *et al*, 2021). Cell-to-cell spread of virus may be less sensitive to neutralization by monoclonal antibodies and convalescent plasma than cell-free virus (Jackson *et al*, 2021). It is thus possible that infected syncytial cells facilitate viral spread. Within this context, it is necessary to better understand the fusogenic potential of the SARS-CoV-2 variants that have arisen and will continue to emerge.

We have characterized here the replication, fusogenicity, ACE2 binding and antibody recognition of Alpha and Beta variants and the role of their S-associated mutations. Despite the insights we provide into the S-mediated fusogenicity of the variants, we did not address the conformational changes that the mutations individually or in combination may elicit. We further show that Alpha, Beta and Delta spike proteins more efficiently bind to ACE2 and are more fusogenic than D614G. Which virological and immunological features of the Delta variant explain its higher estimated transmissibility rate than Alpha and other variants at the population level remains an outstanding question.

## Material and Methods

### Plasmids

A codon optimized version of the reference Wuhan SARS-CoV-2 Spike (GenBank: QHD43416.1) was ordered as a synthetic gene (GeneArt, Thermo Fisher Scientific) and was cloned into a phCMV backbone (GeneBank: AJ318514), by replacing the VSV-G gene. The mutations for Alpha and Beta (Fig. 4A) were added in silico to the codon-optimized Wuhan strain and ordered as synthetic genes (GeneArt, Thermo Fisher Scientific) and cloned into the same backbone (Planas *et al*., 2021a). The D614G S-protein was generated by introducing the mutation into the Wuhan reference strain via Q5 Site-directed mutagenesis (NEB). Other individual mutations were subsequently introduced into the D614G S by the same process. Plasmids were sequenced prior to use. The primers used for sequencing and the site-directed mutagenesis are presented in the supplement **(EV Tables 1 and 2)**. pQCXIP-Empty control plasmid, pQCXIP-IFITM1-N-FLAG, pQCXIP-IFITM2-N-FLAG, pQCXIP-IFITM3-N-FLAG were previously described (Buchrieser *et al*, 2019). pQCXIP-BSR-GFP11 and pQCXIP-GFP1-10 were from Yutaka Hata ((Kodaka *et al*, 2015); Addgene plasmid #68716; http://n2t.net/addgene:68716; RRID: Addgene_68716 and Addgene plasmid #68715; http://n2t.net/addgene:68715; RRID: Addgene_68715). pcDNA3.1-hACE2 was from Hyeryun Choe ((Li *et al*, 2003); Addgene plasmid # 1786; http://n2t.net/addgene:1786; RRID: Addgene_1786). pCSDest-TMPRSS2 was from Roger Reeves ((Edie *et al*, 2018); Addgene plasmid # 53887; http://n2t.net/addgene:53887; RRID: Addgene_53887).

### Cells

Vero E6, HEK293T, U2OS, Caco2/TC7, Calu3 were cultured in DMEM with 10% Fetal Bovine Serum (FBS) and 1% Penicillin/Streptomycin (PS). Vero and 293T GFP-split cells transduced cells with pQCXIP were cultured with 4ug/ml and 1 ug/ml of puromycin (InvivoGen), respectively. U2OS GFP-split cells transduced with pLenti6 were cultured in 1ug/ml puromycin and 10 ug/ml blasticidin (InvivoGen). The MucilAir^TM^ primary human bronchial epithelial model was previously described (Robinot *et al*., 2021). All cells lines were either purchased from ATCC or were kind donations from members of the Institut Pasteur and were routinely screened for mycoplasma.

### Viruses

The Wuhan SARS-CoV-2 strain (BetaCoV/France/IDF0372/2020) and the D614G strain (hCoV-19/France/GE1973/2020) was supplied by Dr. S. van der Werf of the National Reference Centre for Respiratory Viruses (Institut Pasteur, Paris, France). The D614G viral strain was sourced through the European Virus Archive goes Global (EVAg) platform, which is funded by the European Union’s Horizon 2020 research and innovation program under grant agreement 653316. The Alpha strain was isolated in Tours, France, from an individual who returned from the United Kingdom. The Beta strain (CNRT 202100078) originated from an individual in Creteil, France). Informed consent was provided by the individuals for use of their biological materials. The viruses were isolated from nasal swabs on Vero cells and further amplified one or two passages on Vero cells. The viruses were sequenced directly from the nasal squabs and again upon passaging. Titration of Viral stocks was performed by 50% tissue culture infectious dose (TCID50).

### Viral Release

For quantification of extracellular viral RNA, supernatants were diluted and heat-inactivated for 20min at 80°C. qRT-PCR was performed from 1µL of template RNA in a final volume of 5 μL per reaction in 384-well plates using the Luna Universal Probe One-Step RT-qPCR Kit (New England Biolabs) with SARS-CoV-2 N-specific primers (**EV Table 1**) on a QuantStudio 6 Flex thermocycler (Applied Biosystems). Standard curve was performed in parallel using purified SARS-CoV-2 viral RNA. Infectious virus release was assessed by harvesting supernatant at each time point and preforming a TCID50 assay using Vero cells.

### GFP-Split Fusion assay

For cell-cell fusion assays, Vero, U2OS-ACE2, or 293T cell lines stably expressing GFP1-10 and GFP11 were co-cultured at a 1:1 ration (3×10^4^, 2×10^4^ and 7×10^4^ cells/well total, respectively) were transfected in suspension with a total of 100ng of DNA with Lipofectamine 2000 (Thermo) in a 96 well plate (uClear, #655090). 10 ng of phCMV-SARS-CoV2-S and/or 25 ng of pCDNA3.1-hACE2, 25 ng of pCSDest-TMPRSS2, and 40 ng of pQCXIP-IFITM were used and adjusted to 100 ng DNA with pQCXIP-Empty (control plasmid). At 20 h post-transfection images covering 80-90% of the well surface, were acquired per well on an Opera Phenix High-Content Screening System (PerkinElmer). The GFP area and the number of nuclei was quantified on Harmony High-Content Imaging and Analysis Software (Fig EV2B). For infection, cells were plated at the aforementioned concentrations and infected the next day with a range of MOIs and fixed at 20h (U2OS-ACE) or 48h (Vero) post-infection with 4% paraformaldehyde for 30mins. For video microscopy experiments, Vero GFP split cells (mixed 1:1) were transfected in suspension with 50ng of phCMV-SARS-CoV2-S and 450ng of pQCXIP-Empty for 30mins at 37°C. Cells were washed twice and then seeded at a confluency of 2×10^5^ cells per quadrant in a u-Dish 35mm Quad dish (ibidi-#80416). Cells were allowed to settle, and fluorescence images were taken at 37°C every 10min up to 24h using a Nikon BioStation IMQ, with three fields for each condition. Fusion defined as percent of GFP pixels was calculated with ImageJ.

### Flow cytometry

For ACE2 binding, 293T cells transfected with S proteins for 24h were stained with soluble biotinylated ACE2 diluted in MACS Buffer at indicated concentrations (from 60 to 0.01 µg/mL) for 30 min at 4°C. The cells were then washed twice with PBS and then incubated with Alexa Fluor 647-conjugated-streptavidin (Thermo Fisher Scientific, 1:400) for 30 min at 4°C. Finally, the cells were washed twice with PBS and then fixed with 4% paraformaldehyde. The results were acquired using an Attune Nxt Flow Cytometer (Life Technologies). Transfection efficiency was assessed by staining with pan-SARS-CoV-2 human mAb129. Antibody binding to S proteins was assessed via s analogous protocol where transfected 293T cells were first stained with either human mAb10 (pan-coronavirus anti-S2), mAb102 and mAb129 (pan-SARS-CoV-2), mAb48 and mAb98 (SARS-CoV-2 anti-RBD), and mAb71 (SARS-CoV-2 anti-NTD) at 1 µg/mL. The antibodies were derived from convalescent individuals by the Mouquet lab at the Institut Pasteur. mAb10 was generated during the early stages of the epidemic from a patient infected with the Wuhan strain and thus has a higher affinity for the Wuhan spike (Planas *et al*., 2021a). For viral replication, infected cells were fixed at each time with 4% paraformaldehyde for 30 mins. The cells were stained in the same manner described above with anti-spike mAb102 and secondary Alexa Fluor 647 (1:500) in MACS buffer containing 0.05% saponin. The gating strategy to determine spike positive cells is represented in the supplement (Fig EV1B).

### Western Blot

Cells were lysed in TXNE buffer (1% Triton X-100, 50 mM Tris–HCl (pH 7.4), 150 mM NaCl, 5 mM EDTA, protease inhibitors) for 30 min on ice. Equal amounts (20–50 μg) of cell lysates were analyzed by Western blot. The following antibodies were diluted in WB-buffer (PBS, 1% BSA, 0.05% Tween, 0.01% Na Azide): rabbit anti-human TMPRSS2 (Atlas antibodies cat# HPA035787, 1:1,000), rabbit anti-human actin (Sigma cat#A2066, 1:2,000), and human anti-S Serum derived from a convalescent individual (1:1000). Species-specific secondary DyLight-coupled antibodies were used (diluted 1:10,000 in WB-buffer) and proteins revealed using a Licor Imager. Images were quantified and processed using Image Studio Lite software.

### Statistical analysis

Flow cytometry data was analyzed with FlowJo v10 software (Tristar). Calculations were all performed with Microsoft Excel 365. GraphPad Prism 9 was used to generate figures and for statistical analysis. Statistical significance between different conditions was calculated using the tests indicated in the corresponding figure legends.

## Acknowledgments

We thank members of the Virus and Immunity Unit for helpful discussions and Dr. Nicoletta Casartelli for her critical reading of the manuscript. Nathalie Aulner and the UtechS Photonic BioImaging (UPBI) core facility (Institut Pasteur), a member of the France BioImaging network, for image acquisition and analysis support. Work in OS lab is funded by Institut Pasteur, Urgence COVID-19 Fundraising Campaign of Institut Pasteur, ANRS, the Vaccine Research Institute (ANR-10-LABX-77), Fondation Pour la Recherche Médicale (FRM), Labex IBEID (ANR-10-LABX62-IBEID), ANR/FRM Flash Covid PROTEO-SARS-CoV-2 and IDISCOVR. Work in UPBI is funded by grant ANR-10-INSB-04-01 and Région Ile-de-France program DIM1-Health. MMR and MZ are supported by the Pasteur-Paris University (PPU) International Doctoral Program. DP is supported by the Vaccine Research Institute. LG is supported by the French Ministry of Higher Education, Research and Innovation. EB is supported by the Medecine-Sciences ENS-PSL Program. HM lab is funded by the Institut Pasteur, the Milieu Intérieur Program (ANR-10-LABX-69-01), the INSERM, REACTing, EU (RECOVER) and Fondation de France (#00106077) grants. The funders of this study had no role in study design, data collection, analysis, interpretation, or the writing of the article.

## Author contributions

Experimental strategy and design: MMR, JB, MH, LG, RR, LC, OS.

Experimentation: MMR, JB, MH, EB, RR, NS, LG, FGB, FP, RR, JD, SG, AB.

Vital materials and expert advice: CP and HM.

Data processing and figure generation: MMR.

Manuscript writing and editing MMR, JB, OS.

Supervision: JB and OS.

All authors reviewed and approved the manuscript,

---

**Figure EV1.**
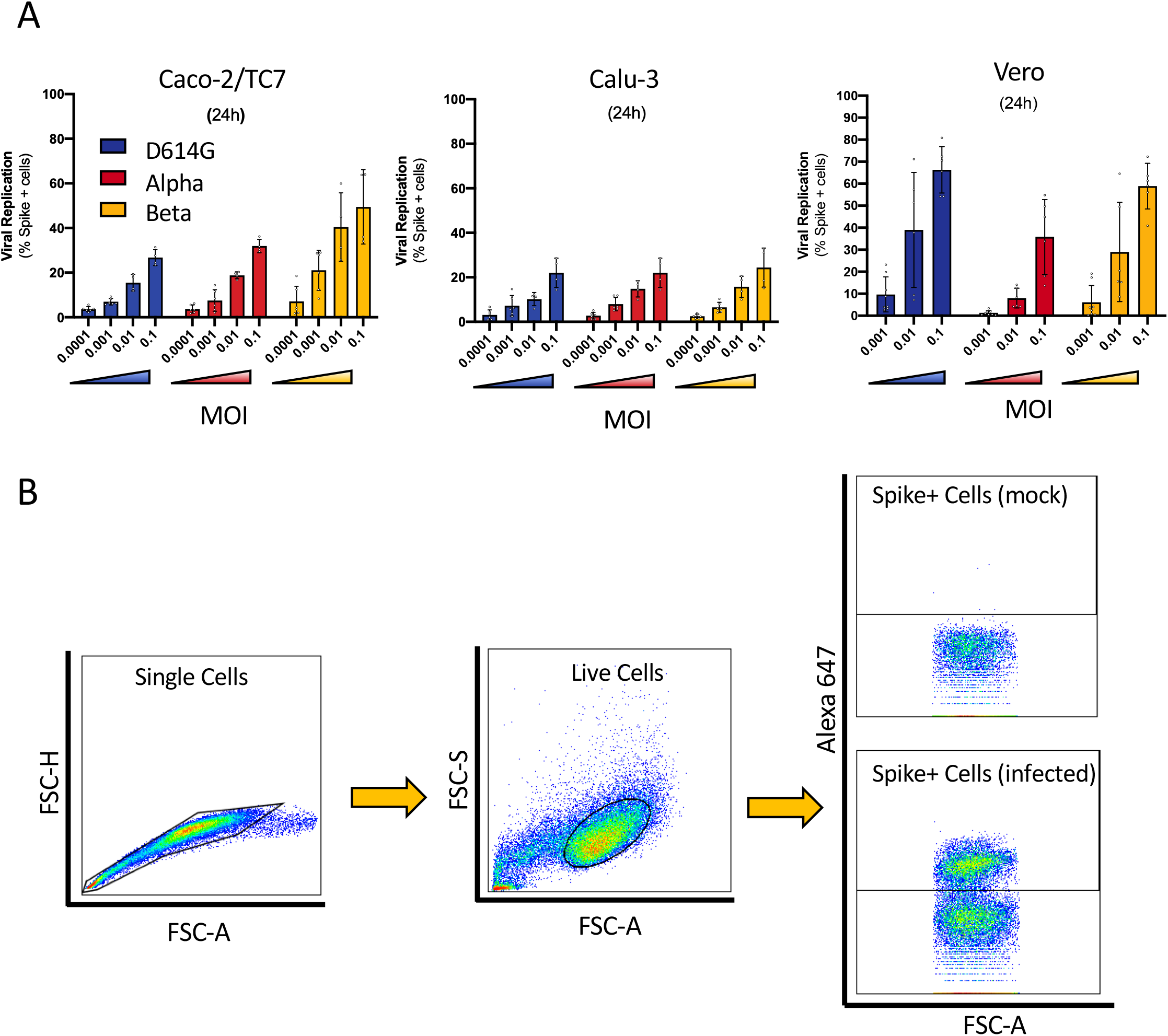
Assessment of viral replication by flow cytometry in cell lines. **(A)** Caco-2/TC7 (left), Calu-3 (middle), and Vero cells (right) were infected at the indicated MOIs with SARS-CoV-2 variants for 24h. The number of spike protein positive cells was determined by flow cytometry upon staining with human pan-SARS-CoV-2 mAb102. Only MOIs that were not saturating were used to generate replication curves for each cell line studied. **(B)** Representative image of gating strategy used for flow cytometry to determine spike positive cells.

**Figure EV2.**
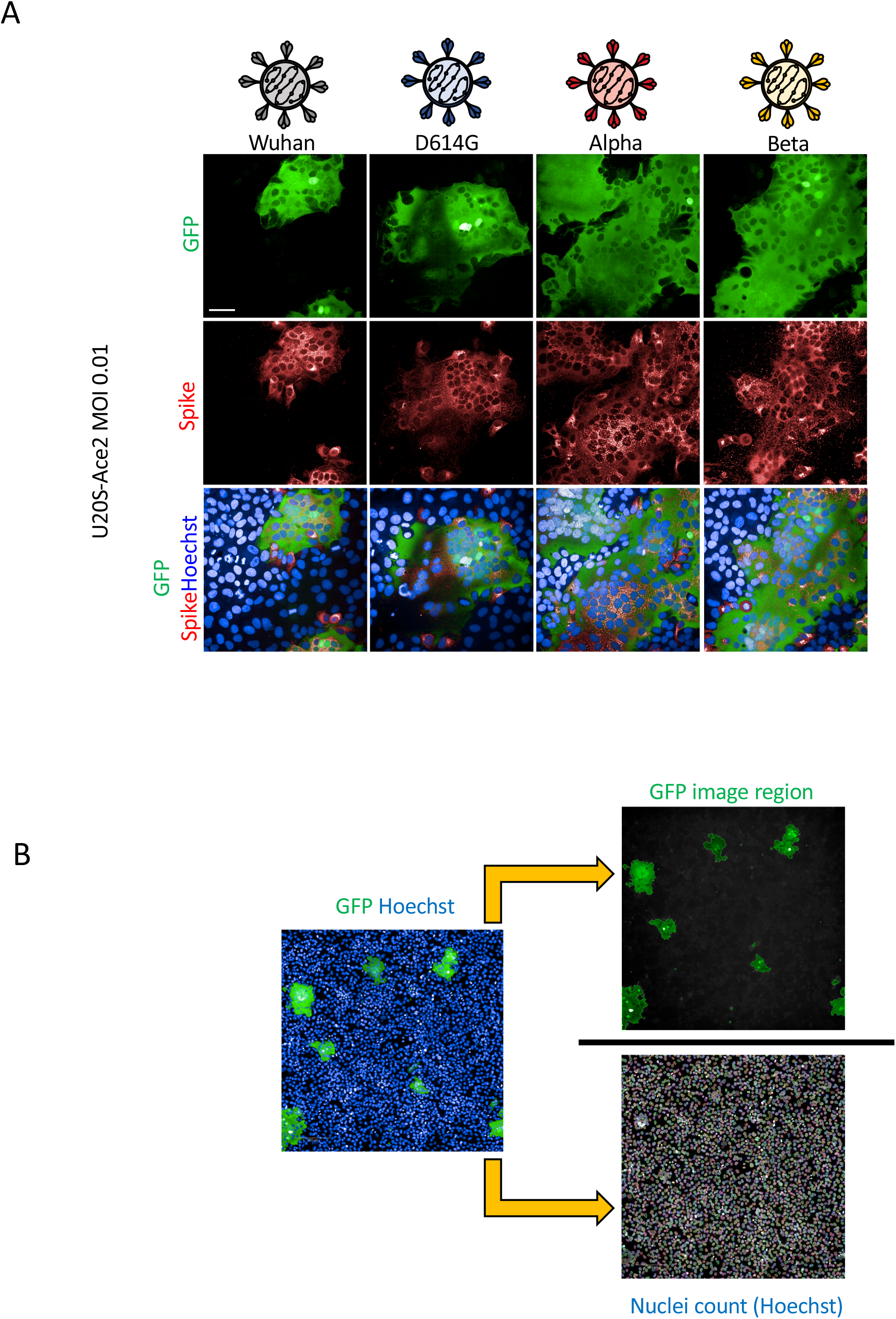
Qualitative and quantitative assessment of syncytia formation. **(A)** U2OS-ACE2 GFP-split cells were infected at MOI 0.01 with the Wuhan, D614G, Alpha and Beta strains for 20h. Cells were stained for spike protein with the human pan-SARS-CoV-2 102 mAb and Alex647 fluorescent secondary antibody. Representative confocal images of the variant induced syncytia formation: GFP-Split (Green), Spike (red) and Hoechst (Blue). **(B)** Quantification method for syncytia formation using the Opera Phenix high content imager and harmony software: Total syncytia area (GFP area) is normalized for cell number upon quantifying the number of nuclei (Hoechst). Scale bars: 50 µm

**Figure EV3.**
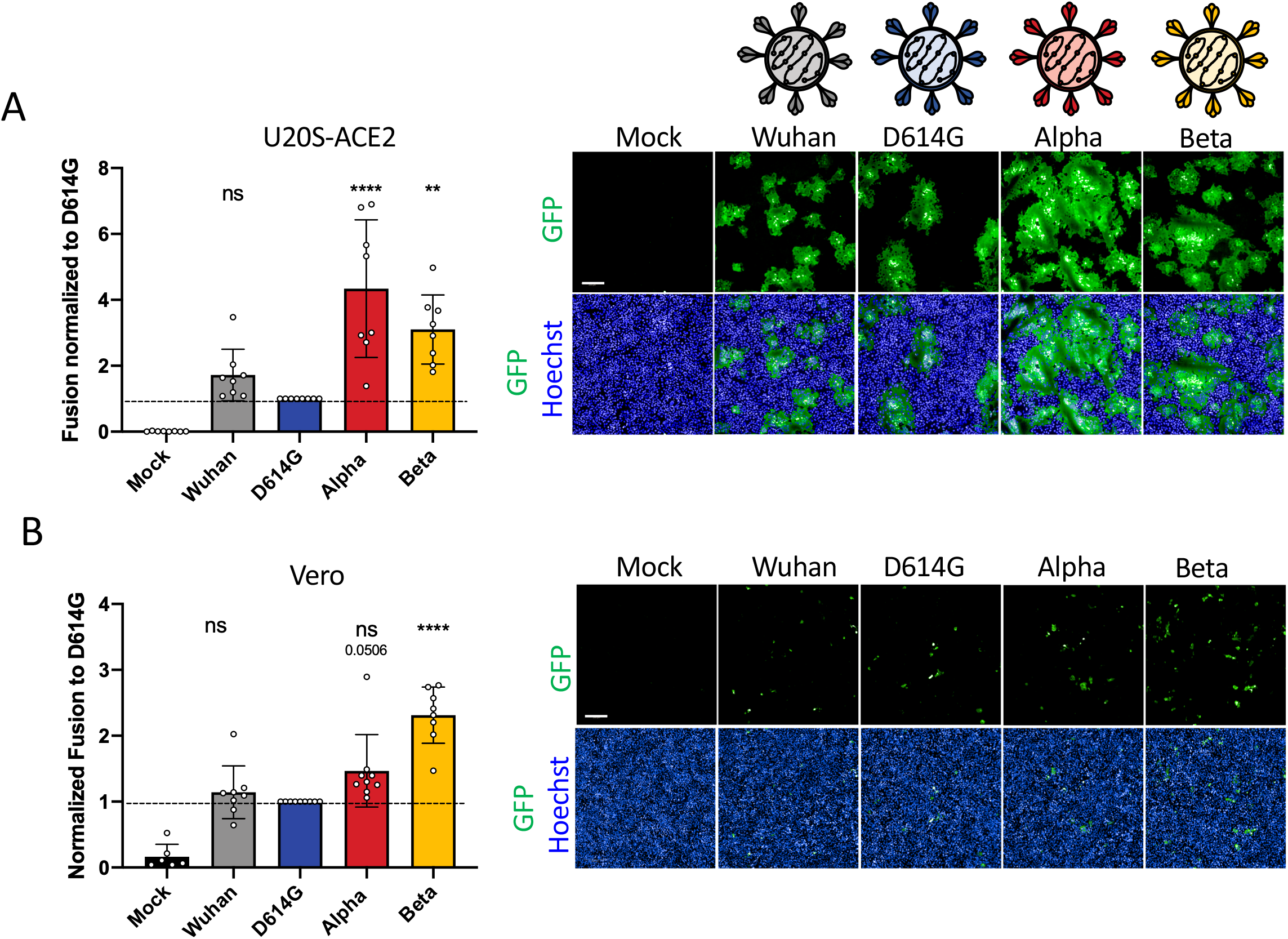
Syncytia formation by SARS-CoV-2 variants. **(A) Left Panel:** Fusion normalized to D614G for U2OS-ACE2 20h post infection at MOI 0.01. **Right Panel:** Representative images of U2OS-ACE2 20h post infection, GFP (Green) and Hoechst (Blue). Top and bottom are the same images with and without Hoechst channel. **(B) Left Panel:** Quantified fusion of Vero cells infected at MOI 0.1. **Right Panel:** Representative images of Vero cells 48h post infection. Scale bars: 200 µm. Data are mean ± SD of at least 3 independent experiments. Statistical analysis: One-way ANOVA compared to D614G reference, ns: non-significant, *P < 0. 05, **P < 0.01, ***P < 0.001, ****P < 0.0001.

**Figure EV4.**
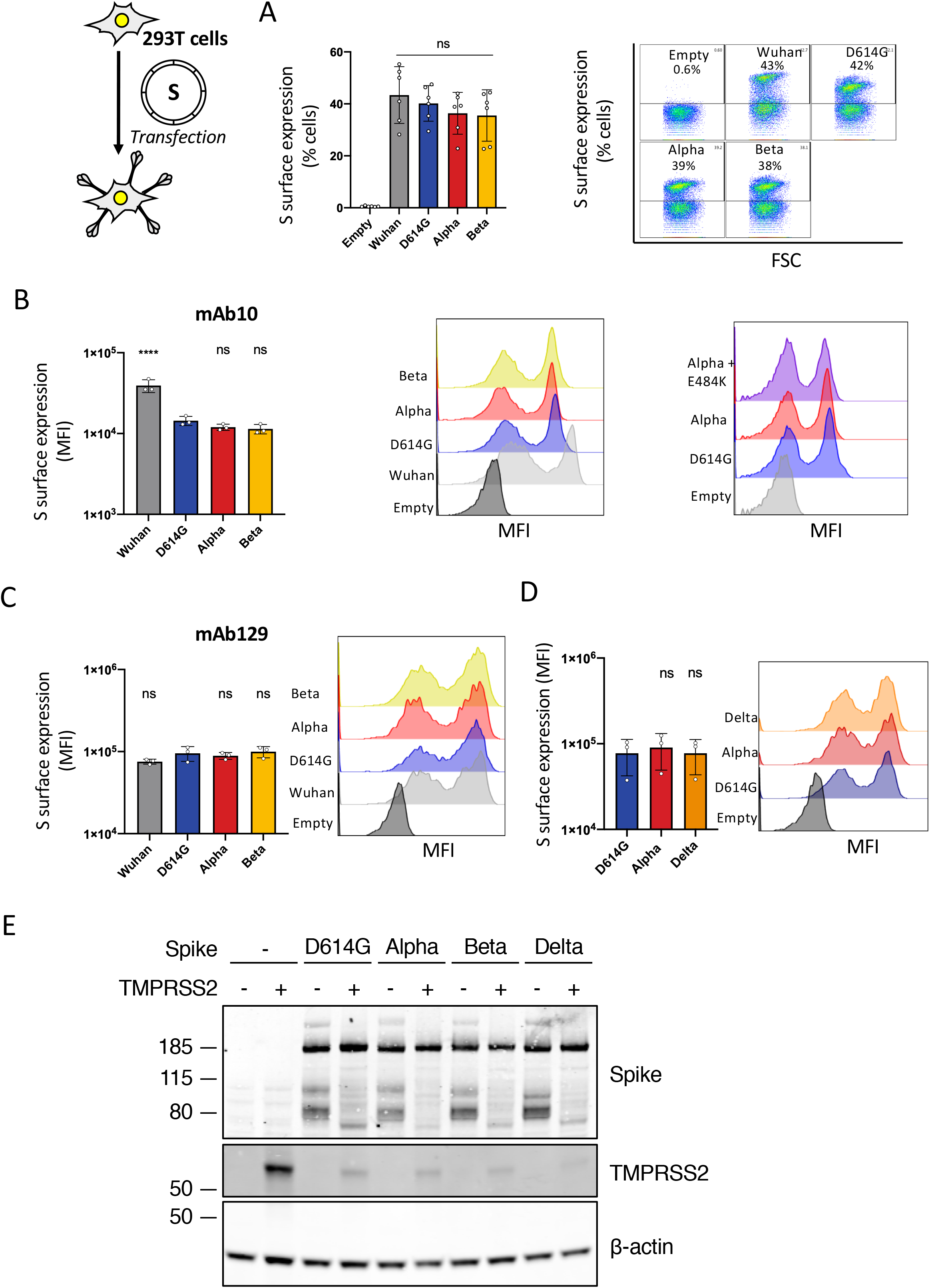
SARS-CoV-2 variant S proteins are expressed equally at the cell surface. 293T cells were transfected with variant S proteins for 20h and stained with human pan-coronavirus mAb10 without permeabilization. 293T cells were chosen because they lack ACE2 and do not fuse upon S transfection; this makes them suitable for single cell flow cytometry **(A) Left Panel:** Quantification of percent of cells expressing each spike at the surface. **Right Panel:** Representative FACs plots. **(B)** Quantification of median florescent intensity (MFI) of variant spikes at the cell surface and representative histograms of MFI of the Wuhan, D614G, Alpha, Beta, and Alpha + E484K variants spikes using mAb10. **(C)** Quantification of median florescent intensity (MFI) of variant spikes at the cell surface and representative histograms of MFI of the Wuhan, D614G, Alpha, Beta, and Alpha + E484K variants spikes using mAb129. **(D)** Quantification of median florescent intensity (MFI) of variant spikes at the cell surface and representative histograms of MFI of the Delta variant compared to the Alpha and D614G using mAb129. **(E)** Comparison of the impact of TMPRSS2 on variant S protein processing measured by western blot. Plasmids encoding for S protein were co-transfected with or without plasmids expressing TMPRSS2 in 293T cells for 24h. Representative image of 2 experiments. Flow cytometry data are mean ± SD of at least 3 independent experiments. Statistical analysis: One-way ANOVA compared to D614G reference, ns: non-significant, *P < 0. 05, **P < 0.01, ***P < 0.001, ****P < 0.0001.

**Figure EV5.**
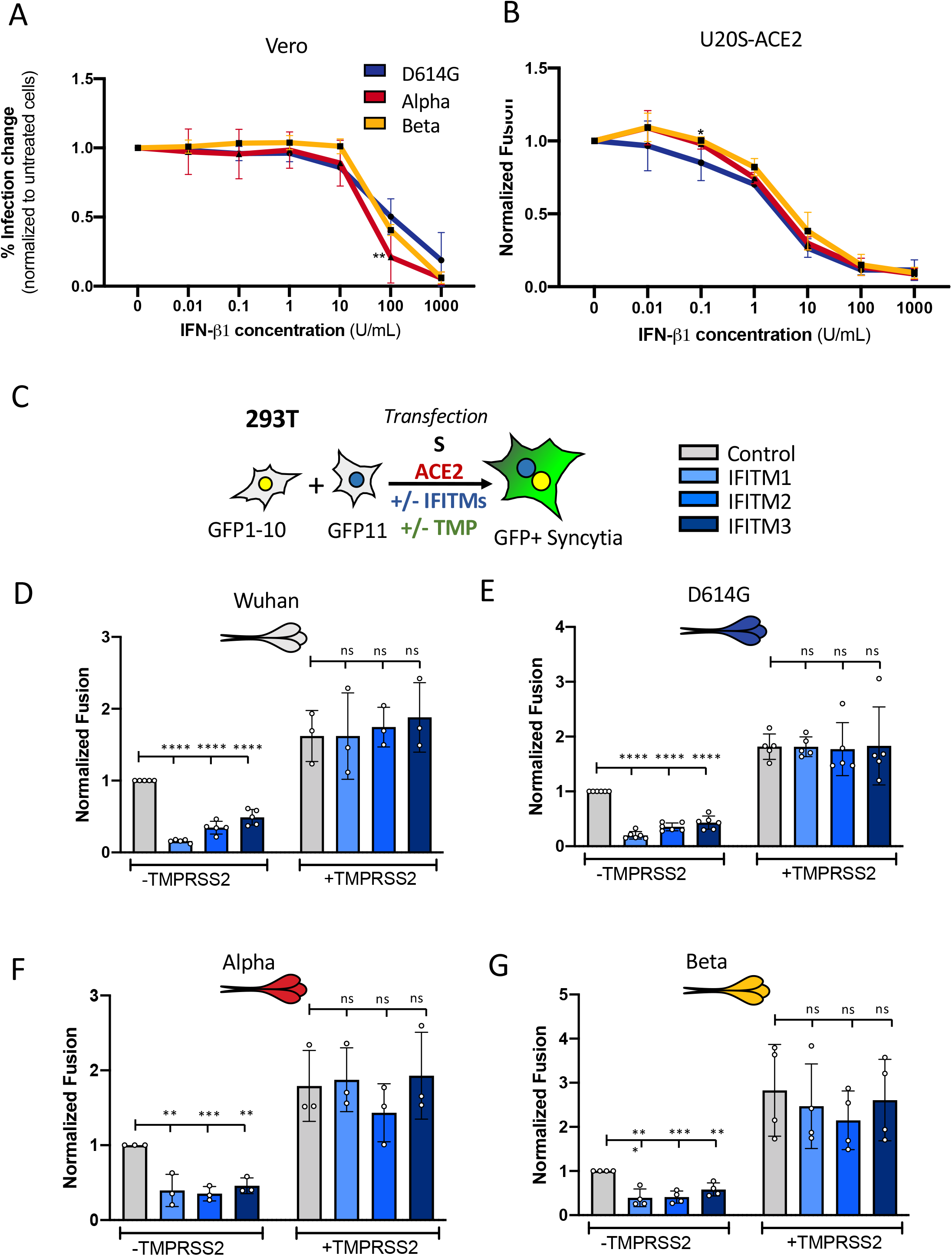
Impact of IFN-β1 and IFITMs on SARS-CoV-2 variant replication and S protein mediated cell-cell fusion. **(A)** Vero cells were pre-treated for 2h with a serial dilution of IFN-β1 prior to infection with the SARS-CoV-2 variants. Infected cells were maintained in media containing IFN-β1 and analyzed by flow cytometry 48h post-infection to determine relative infection change. **(B)** U20S-ACE2 GFP split cells were pre-treated for 2h with a serial dilution of IFN-β1 prior to infection with the SARS-CoV-2 variants. Infected cells were maintained in media containing IFN-β1 and relative inhibition of syncytia formation 20h post-infection was determined via GFP signal. **(C)** A co-culture of 293T GFP-Split cells were transfected with combination of S, control, ACE2, TMPRSS2 and IFITM plasmids and then imaged 18h post-transfection. Effect of IFITMs and TMPRSS2 on the cell-cell fusion induced by different spike proteins **(D)** Wuhan **(E)** D614G **(F)** Alpha **(G)** Beta. Data are mean ± SD of at least 3 independent experiments. Statistical analysis: One-way ANOVA compared to D614G reference or control plasmid transfection, ns: non-significant, *P < 0. 05, **P < 0.01, ***P < 0.001, ****P < 0.0001.

**Figure EV6.**
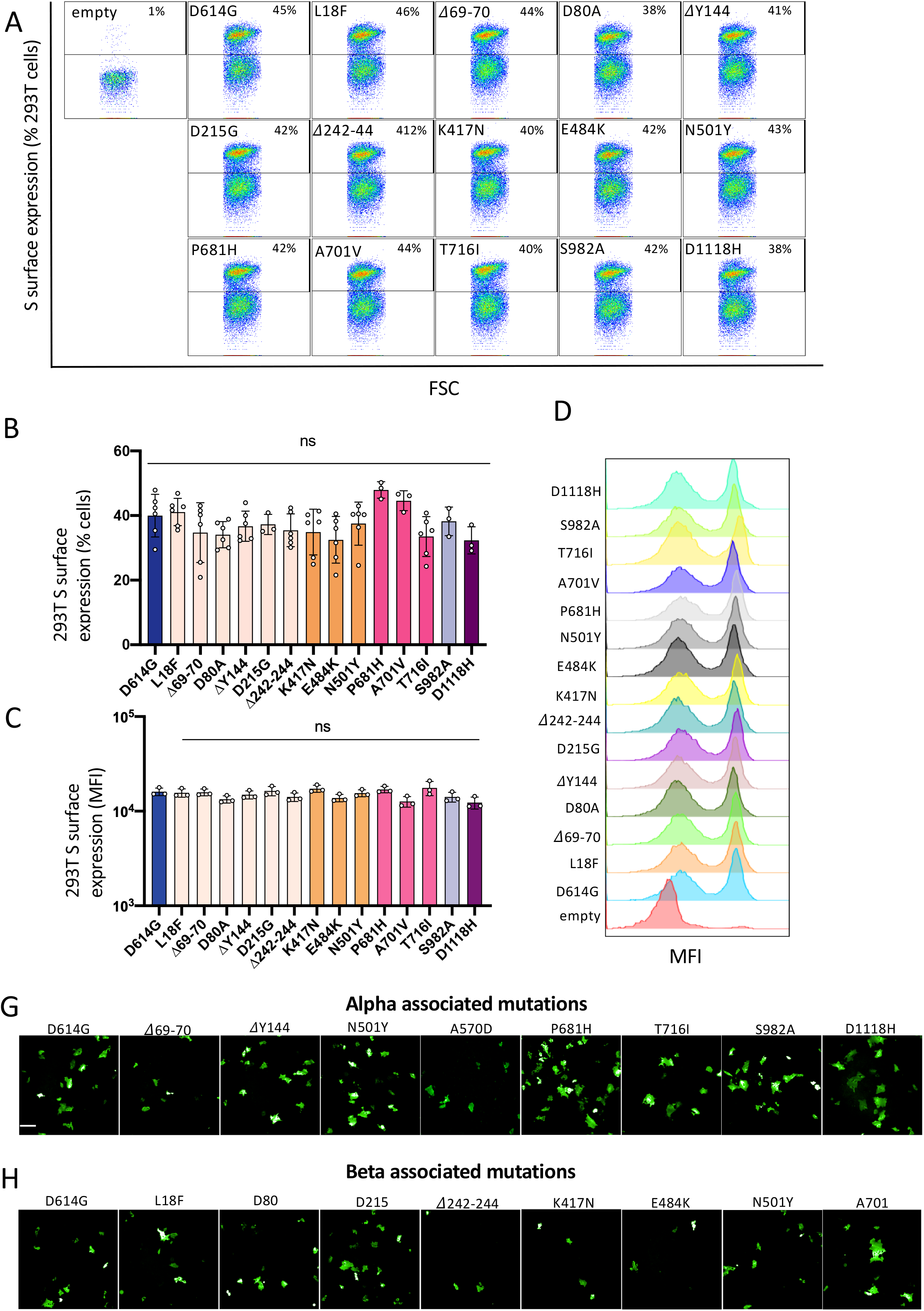
SARS-CoV-2 variant S protein associated mutations are expressed equally at the cell surface. 293T cells were transfected with S proteins with each of the variant associated mutations for 18h and stained with human pan-coronavirus mAb10 without permeabilization. **(A)** Representative FACs plots of percent of cells expressing each mutant spike at the surface. **(B)** Quantification of percent of cells expressing each spike at the surface. **(C)** Quantification of median florescent intensity (MFI) of the mutant spikes at the cell surface. **(D)** Representative histograms of MFI of each mutant spike. **(E)** Representative images of Vero GFP split cells 20h after transfection with each Alpha variant associated mutant spike, GFP-Split (Green). **(F)** Representative images of Vero GFP split cells 20h after transfection with each Beta variant associated mutant spike. Statistical analysis: One-way ANOVA compared to D614G reference, ns: non-significant, *P < 0. 05, **P < 0.01, ***P < 0.001, ****P < 0.0001.

**Figure EV7.**
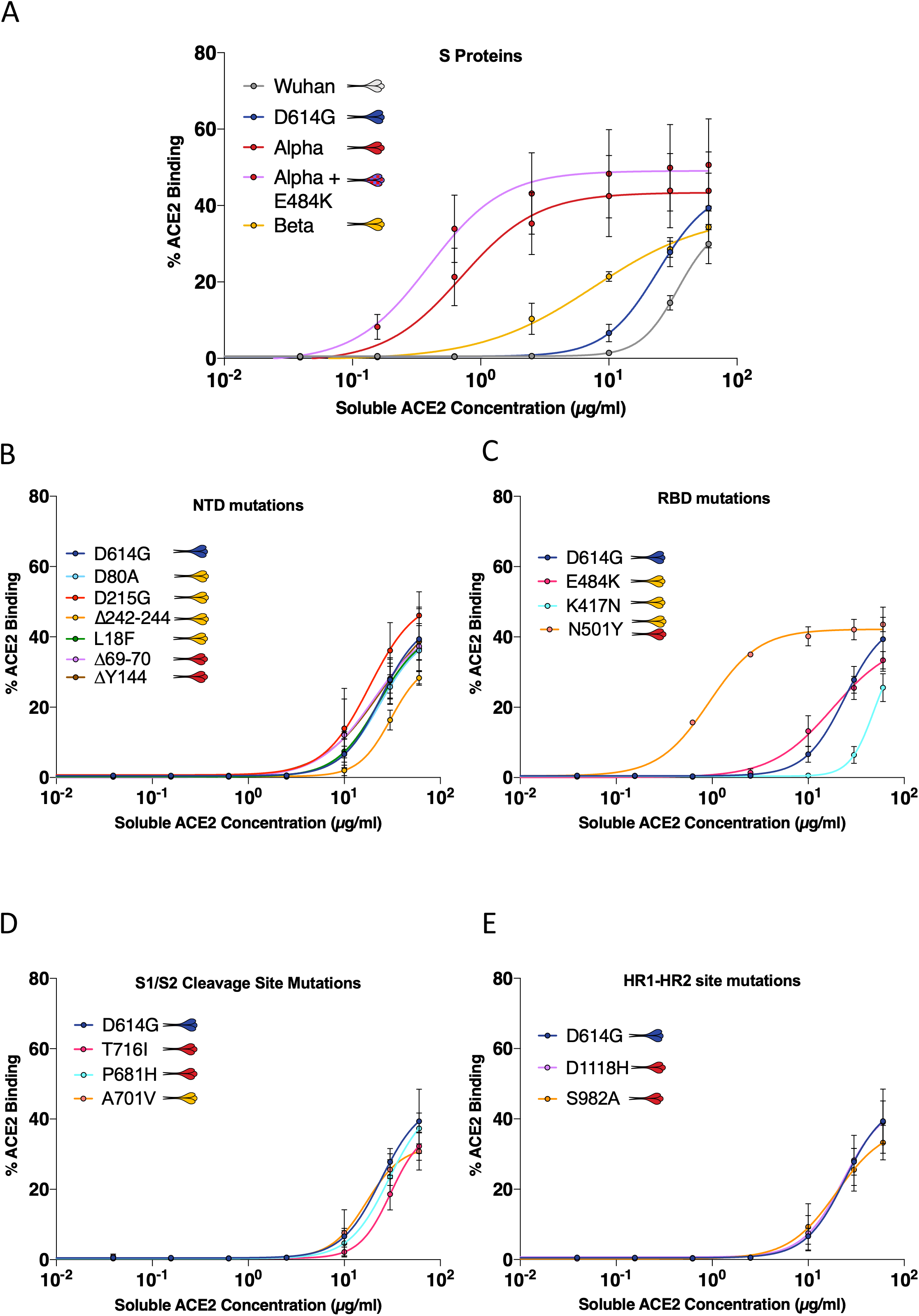
ACE2 binding curves to SARS-CoV-2 variant S proteins and associated mutations. 293T cells were transfected with variant or mutant spike proteins for 24h and stained with a serial dilution of soluble biotinylated ACE2 and revealed by fluorescent streptavidin before analysis by flow cytometry. **(A)** ACE2 binding dilution curves of each variant spike. **(B)** ACE2 binding dilution curves of each variant associated mutation located in spike n-terminal domain (NTD) **(C)** receptor binding domain (RBD) **(D)** S1/S2 cleavage site **(E)** heptad repeat 1-2 site (HR1-HR2). Data are mean ± SD of 3 independent experiments.

**Movie EV1: Alpha (B.1.1.7) and Beta (B.1.351) S proteins induce more robust syncytia formation than D614G or Wuhan**

A co-culture of Vero GFP-split cells were transfected with plasmids expressing each of the variant spike proteins and imaged by video microscopy at a rate of 6 images per hour for 24h. GFP signal is superimposed over BF images. The white border represents the GFP area calculated by the ImageJ macro in order to quantify fusion. **Top left:** Wuhan S **Top right:** D614G S **Bottom Left:** Alpha (B.1.1.7) S **Bottom right:** Beta (B.1.351) S. One representative field for each condition is shown.

